# Social signaling via bioluminescent blinks drives schooling behavior in the flashlight fish *Anomalops katoptron*

**DOI:** 10.1101/2020.06.16.143073

**Authors:** Peter Jägers, Louisa Wagner, Robin Schütz, Maximilian Mucke, Budiono Senen, Gino V. Limmon, Stefan Herlitze, Jens Hellinger

## Abstract

The bioluminescent flashlight fish *Anomalops katoptron* live in schools of several hundred specimens. To understand how flashlight fish, integrate bioluminescent signaling into their schooling behavior, we analyzed movement profiles and blink frequencies. Isolated specimen of *A. katoptron* show a high motivation to align with fixed or moving artificial light organs. Depending on presented frequencies *A. katoptron* responds with a reduction in swimming speed and its own blink frequency. Higher presented blink frequencies reduce the nearest neighbor distance. In the natural environment *A. katoptron* is changing its blink frequencies and nearest neighbor distance in a context specific manner. Blink frequencies are increased from day to night and during avoidance behavior, while nearest neighbor distance is decreased with increasing blink frequencies. *A. katoptron* changes its blink frequencies by modifying light organ occlusion. Our results suggest that visually transmitted information via specific blink patterns determine intraspecific communication and group cohesion in schooling *A. katoptron*.

## Introduction

Bioluminescence is a widespread phenomenon in ocean-dwelling organisms including a broad phylogenetic distribution in marine fish ^1^. In ray finned fish bioluminescence evolved independently at least 27 times ^2^. In vertebrates only fish possess the ability to emit light via own photophores, bioluminescent bacteria hosted within specialized light organs or kleptoproteins acquired from prey ^3^.

Numerous functions of bioluminescence have been described and suggested such as counterillumination ^4,5^, mate attraction ^6^, prey attraction ^7^ and prey illumination in flashlight fish (Anomalopidae) ^8^. All members of the family Anomalopidae such as Photoblepharon and Anomalops are characterized by bean-shaped, subocular light organs ^9,10^. In *Photoblepharon steinitzi* three distinct functions in bioluminescent signaling like finding prey, intraspecific communication and confusing predators via a “blink and run-pattern” have been proposed ^11,12^. *Photoblepharon* reside solitary- or pairwise in territories (e.g. reef caves) while *Anomalops katoptron* (Anomalopidae) occur in large, moving schools during moonless nights ^8,13^.

The extrinsic, continuous bioluminescent light in *A. katoptron* is produced by symbiotic bioluminescent bacteria *Candidatus photodesmus katoptron* (Gammaproteobacteria: Vibrionaceae) hosted within subocular light organs. Anomalopid symbionts show a genome reduction like other unrelated, obligate symbiotic bacteria, such as insect endosymbionts. It has been proposed that symbionts of *A. katoptron* are transmitted during an active environmental phase ^14–16^. Symbiotic bacteria are densely packed in numerous tubules that are aligned at right angles to the light-emitting surface of light organs ^8,16,17^. The inner surface of light organs contains two stacks of guanine crystals, which serve as reflector to enhance light emission ^18^. At the anterior edge light organs are attached to suborbital cavities via the rod like “Ligament of Diogenes” which allows a downward rotation. This exposes the dark pigmented back of light organs and disrupts light output. The visual system of *A*. *katoptron* is optimized to detect wavelengths in the frequency range of its own bioluminescent symbionts ^19,20^. Fascinating blink patterns of large schools can be observed on coral reefs in the Indo-Pacific during dark and moonless nights ^13,21^. During the daytime *A. katoptron* hides in crevices, caves or deep water ^8,21^.

In general, groups of fish show various formations ranging from lose aggregations to highly aligned groups moving in synchronized directions ^22,23^. Living in a group can be advantageous in several aspects like lower predation risk, mate choice ^24^, reduced metabolic costs ^25^ and higher probability in detecting food sources ^26^. It has been proposed that a synchronized organization within the school leads to lower vulnerability ^27^. Group size and cohesion play an important role in schooling and can reduce the risks of being preyed through attack abatement ^28^ or confusion of predators ^29^.

The ability to sense intraspecific group members is important to maintain the formation of a school ^30^. Sensory input from vision and lateral lines are integrated to determine attraction or repulsion in moving groups. Partridge & Pitcher suggested that vision is primarily used for maintenance of position and angle between fish while lateral lines monitor swimming speed and direction of moving neighbors ^31^. The school formation is situation-dependent and can be interpreted as an integration of surrounding ecological factors. For example higher predation regimes force shoaling groups of *Poecilia reticulata* (Poecillidae) to form denser aggregations with closer nearest neighbor distance ^27,32^. Collective behavior has been recently analyzed with computer models and/or robotic dummies revealing strong correlation between decision rules of individuals driving group behavior ^33–36^.

Providing information to conspecifics is an important feature to maintain the functionality of a dynamic group and can be observed on inter-individual and/or group level ^35^. Many different ways of intraspecific communication are described within fish just as mutual allocation in the weakly electric fish *Mormyrus rume proboscirostris* (Mormyridae) via electrocommunication that leads to social attention ^37^ or startle response as a reaction on moving neighbors in *Clupea harengus* (Clupeidae) ^38^.

As nocturnal animals live under visual restriction, bioluminescent signaling can become an additional source of information ^7^ e.g. in orientation towards conspecifics shown in ostracodes (Cypridinidae) ^39,40^, dragonfish (Stomiidae) ^41,42^ or pony fish (Leiognathidae) ^43^. For *Gazza minuta* (Leiognathidae) discrete projected luminescent flashes have been described. Possible functions are spacing between foraging individuals, keeping the school together or reproductive activities each represented in different flash patterns ^44^.

It has been shown that *A. katoptron* uses its light organs to actively localize food. During feeding the light organs reveal a prolonged exposure and shorter occlusion time resulting in decreased blink frequencies ^8^. In addition, it has been described that the light organs play a role in orientation towards conspecifics in schooling behavior of *A. katoptron* ^13^.

In this study we investigated how *A. katoptron* behaviorally responds to different artificial light stimuli and if these behavioral responses can be compared to a context-dependent blinking behavior observed in the natural environment at the Banda Sea. We found that *A. katoptron* is attracted by blue green light (500 nm) in a blink frequency and light intensity dependent manner. The fish responds with an adjustment of its own blink frequencies, where the light organ occlusion, but not the exposure time is adjusted. Higher blink frequencies are correlated with closer nearest neighbor distance leading to a higher group cohesion. Thus, our study shows for the first time that the blink frequencies of the bioluminescent light of the flashlight fish *A. katoptron* is used for a context dependent, intraspecific communication.

## Results

To investigate how bioluminescent signaling emitted by the light organs of the splitfin flashlight fish *Anomalops katoptron* is used for intraspecific communication, we investigated the behavioral responses of isolated flashlight fish to artificial light pulses in the laboratory. It has been suggested that *A. katoptron* in its natural environment reveal a schooling behavior. To investigate if and how *A. katoptron* reacts to different light signals we isolated *A. katoptron* in an experimental tank (Fig. 1A, 1B). In the middle of the tank we introduced a light emitting dummy and defined two areas, where we analyzed the duration of how long the fish spend in this area, i.e. one area close to the dummy (center area) and one area close to the exit area (exit door) of the tank. This exit door is normally open and connects the experimental tank with the housing tank. Without light stimulation individuals stayed for 64.4 ± 5.7 % (Fig. 1A, 1C) of the time in the exit area compared to the center area. Fish were swimming with a swimming speed of 0.19 ± 0.01 m/s (Fig. 1D). Stimulation with artificial light organs caused an orientation towards the fish dummy (Fig. 1B). Isolated specimen spent 79.7 ± 3.9 % of the time in the center around the light emitting dummy (LED timing: 2 Hz, 0.25 s on + 0.25 s off) and reduced their swimming speed. (Fig. 1C and 1D, Video S1).

**Figure 1.**
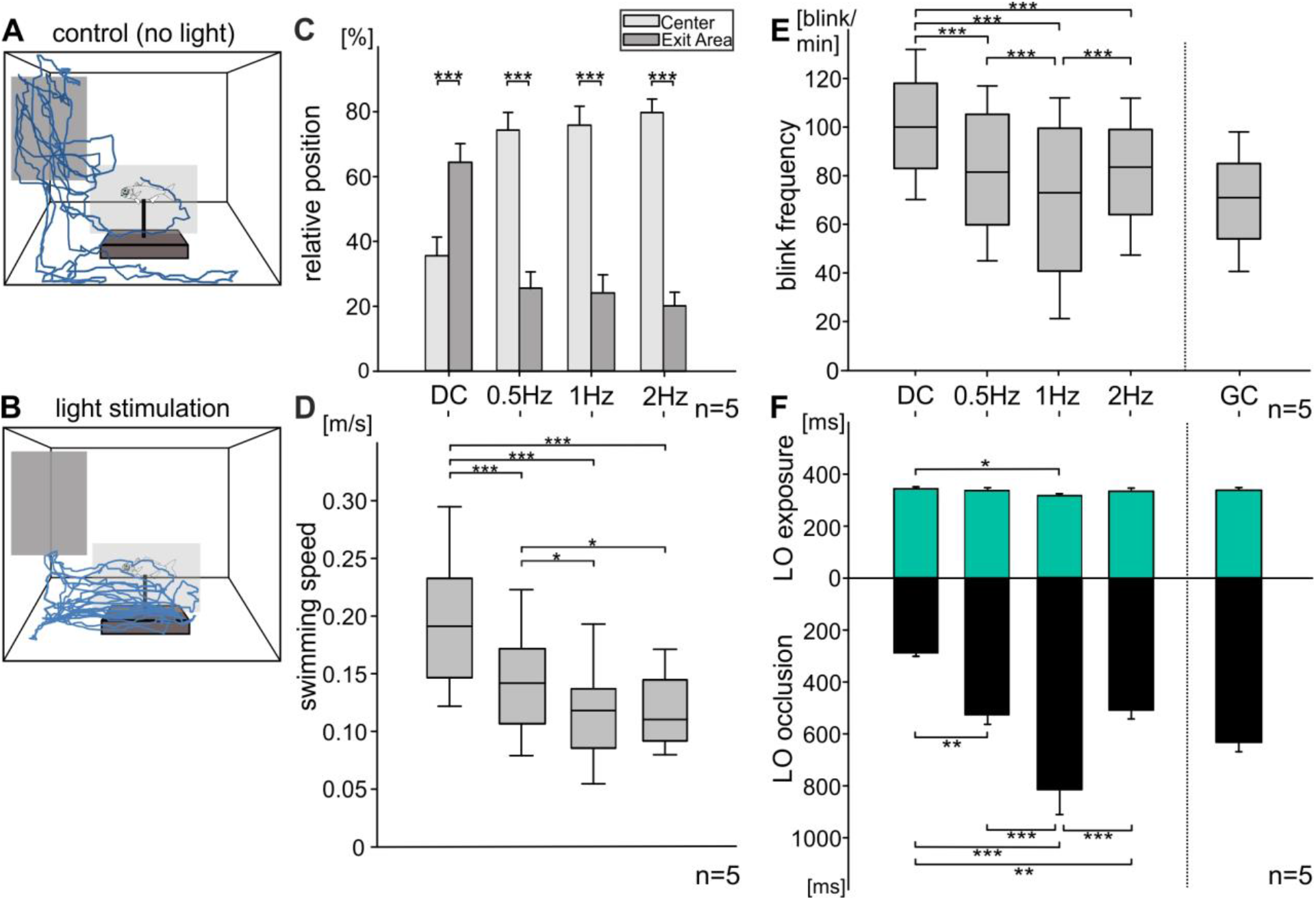
Changes in positioning, swimming speed, blink frequency and light organ occlusion of flashlight fish (*A. katoptron*) induced by a fish dummy equipped with artificial light organs. (A, B) Example of two-dimensional trajectories for (A) control without and (B) 1 Hz stimulation (LED light pulses with equal distributed on- and off times) of isolated flashlight fish (n=5). Boxes indicate the two defined areas of interest, which were analyzed. Dark gray is defined as “Exit Area” and light gray as “Center”, where the dummy with artificial light organs was placed. Each trajectory was traced for 60 s. (C) Relative Positioning (± SEM) of isolated *A. katoptron* (n=5) in areas of interest during four different stimuli (DC, dark control; 0.5, 1, 2 Hz blinking LED with equal distributed LED on- and off-times within artificial light organs). (D) Swimming speed (m/s) of isolated *A. katoptron* (n=5) during four different stimuli (DC; 0.5, 1, 2 Hz blinking LED with equal distributed LED on- and off-times). (E) Blink frequencies of *A. katoptron* (n=5) induced by four stimuli (DC; 0.5, 1, 2 Hz blinking LED with equal distributed LED on- and off-times). Additionally, blink frequencies of a group consisting of five individuals (GC), which is divided by a scattered line, were analyzed. Individuals were tested separately before group experiments. (F) Mean light organ exposure and occlusion time (± SEM) during four different stimuli and a group control (DC, dark control; 0.5, 1, 2 Hz blinking LED within artificial light organs with equal distributed LED on- and off-times; GC, group control). Upper lines refer to stimulation as seen in (E). Greenish bars indicate exposure of light organs and occlusion of light organs is represented by black bars (n=5). DC, dark control; GC, group control. Significance values are reported as *p < 0.05, **p < 0.01, ***p < 0.001. Error bars indicate ± SEM.

Control experiments showed that the shape of the dummy does not have an impact on the behavior of *A. katoptron* (Fig. S1). These findings suggest that light pulses are used for intraspecific communication of *A. katoptron* and that *A. katoptron* is attracted by these light pulses (Fig. 1).

To investigate if the light intensity of light pulses plays a role for intraspecific communication, we determined the emitted light intensity of *A. katoptron’s* light organs. Light emitted by luminous bacteria housed within the light organs of *A. katoptron* had a maximum intensity of 0.27 μW (emission peak at 510 nm wavelength, n=5; Fig. S2). We next applied LED light stimuli (1 Hz, 0.5 s on + 0.5 s off) with light intensities of 0.12, 0.33 and 1.52 μW (at 504 nm wavelength) and found that increasing light intensities resulted in decreased blink frequencies. There were no differences in distances kept to the dummy (around 15.67 ± 0,64 cm). Thus, throughout the experiments we used a LED light, with an intensity at 504 nm wavelength of 0.23 μW (except for the intensity experiments), which is slightly dimmer than the light emitted from the light organ of *A. katoptron*.

To investigate if the blink frequency is important for intraspecific communication, we presented three different blink frequencies (0.5 Hz; 1 Hz & 2 Hz) with equally distributed LED light on- and off-times (Fig. 1C–F). While there was no difference in time spent in the center area (Fig. 1C), there was a frequency-dependent change in swimming speed (Fig. 1D), the blink frequency response (Fig. 1E) along with the exposure and occlusion of the light organs (Fig. 1F). A light stimulation of 0.5 Hz resulted in a swimming speed of 0.146 ± 0.009 m/s, which is faster than the swimming speed determined for 1 Hz and 2 Hz stimulation (1 Hz (0.115 ± 0.008 m/s), 2 Hz (0.119 ± 0.006 m/s), RM ANOVA 0.5 Hz compared to: 1 Hz, p = 0.014; 2 Hz, p = 0.023; Fig. 1D).

We next analyzed the blink frequency responses of *A. katoptron*. We found that during schooling behavior in the tank the average blink frequency of individuals was 1.17 Hz (69.88 ± 1.78 blinks/min), while in isolation the blink frequency is increased to 1.67 Hz (100.39 ± 1.83 blinks/min). At 1 Hz LED light stimulation, the blink frequency of *A. katoptron* was 70.25 ± 2.72 blinks/min and was comparable to the blink frequency within the school (i.e. 1.17 Hz), but is increased to 1.35 Hz for 0.5 Hz and 2 Hz light stimulation.

Next we investigated mean light organ exposure and occlusion for the different experimental light pulse settings. We found that the time individuals expose light organs is around 330 ms, which was comparable throughout the experiments (DC (344 ± 0.005 ms), 0.5 Hz (338 ± 0.004 ms), 1 Hz (317 ± 0.006 ms), 2 Hz (336 ± 0.008 ms); Fig. 1F). In contrast, differences existed in how long the organ is occluded. We found that in isolation the fish decreases its occlusion time to 287 ± 0.01 ms, while during schooling (618 ± 0.069 ms) and in the presence of the light stimuli, light organ occlusion increased (0.5 Hz (528 ± 0.035 ms), 1 Hz (967 ± 0.092 ms), 2 Hz (0.507 ± 0.036 ms); Fig. 1F). These findings suggest that light organ occlusion defines blink frequencies during schooling.

Thus, the findings on blink frequencies related to light organ occlusion, orientation and swimming speed led us to the hypothesis that the timing of light pulses emitted by *A. katoptron* bear information to keep attraction and alignment of *A. katoptron* to its conspecifics.

To investigate this hypothesis, we established a second experimental setup in a circular arena tank, with a light pulse emitting dummy in the middle of the arena (Fig. 2A). We changed the LED off-times between 200 ms and 500 ms with on-times at 300 ms and examined the distance of the individuals towards the artificial light organs of the dummy in the center using heat maps. Without light stimulation individuals were swimming along the wall and avoiding the middle of the arena (Fig. 2C1) with a mean distance of 42.25 ± 0.76 cm to the dummy (Fig. 2B). During light stimulation *A. katoptron* changed its swimming behavior in an off-time dependent manner (Fig. 2C2-3). A 500 ms LED off-time resulted in a closer but still partly decentralized orientation (23.63 ± 0.88 cm) towards the dummy in comparison to the dark control (DC; RM ANOVA: p < 0.001, Fig. 2C2; Video S2). The closest and centralized orientation towards the LED dummy occurred with 200 ms off-time LED stimulation (Fig. 2C3). These findings suggest that light organ occlusion contains information about nearest neighbor distance for *A. katoptron*.

**Figure 2.**
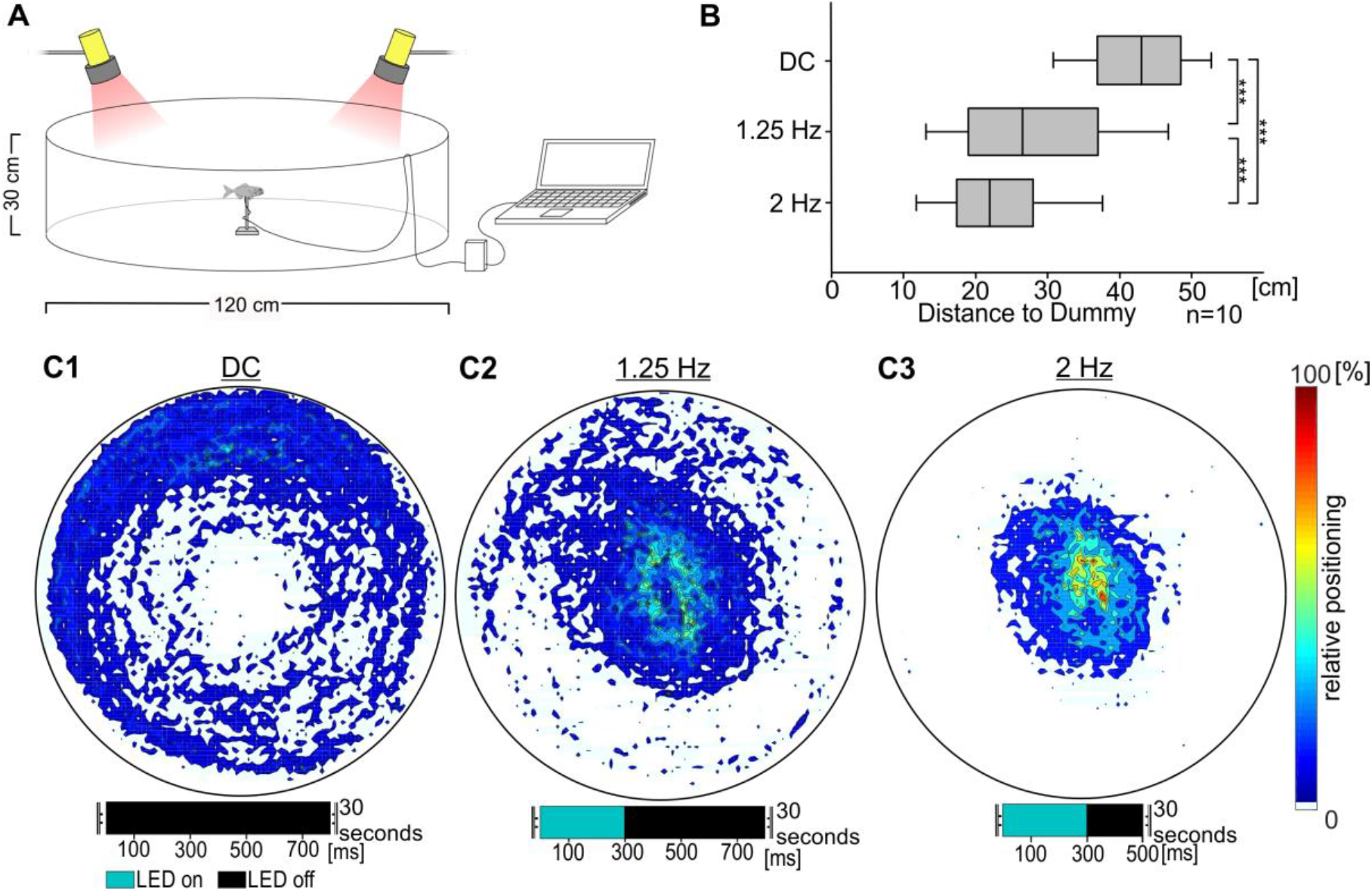
Nearest neighbor distance communicated via artificial light organs within flashlight fish (*A. katoptron*). (A) Experimental setup for the validation of changes in nearest neighbor distance. Artificial light organs of a center placed dummy were emitting different light stimulations (1.25 Hz, LED off-time 500 ms; 2 Hz LED off-time 200 ms) for 30 seconds. An additional control without light stimulation was performed (DC, dark control). In this experiment LED off-timing was adjusted while LED on-time was consistent for 300 ms. (B) Distance between isolated specimen (n=10) and the center placed fish dummy equipped with artificial light organs. (C) Heat Maps indicate relative positioning of *A. katoptron* (n=10) in relation to a light (C2/C3) or no light (C1) emitting dummy. A closer orientation can be observed with shorter intervals (200 ms) between constant light emittance of 300 ms. Without light stimulation individuals show a wall following behavior. Heat Maps are based on all trajectories recorded for each stimulation. DC, dark control. Significance values are reported as: *p < 0.05, **p < 0.01, ***p < 0.001

In the ocean, schools of *A. katoptron* constantly move through the open water, suggesting that individuals recognize/monitor their nearest neighbor to stay aligned. Thus, we next examined if *A. katoptron* would follow a moving light signal. To perform this experiment, we used an experimental setup, in which 13 LEDs arranged in a circular swimming tank separated by an angle of 27.7 ° lit up for 300 ms consecutively clockwise or counterclockwise (Fig. 3A, S3 and Video S3).

**Figure 3.**
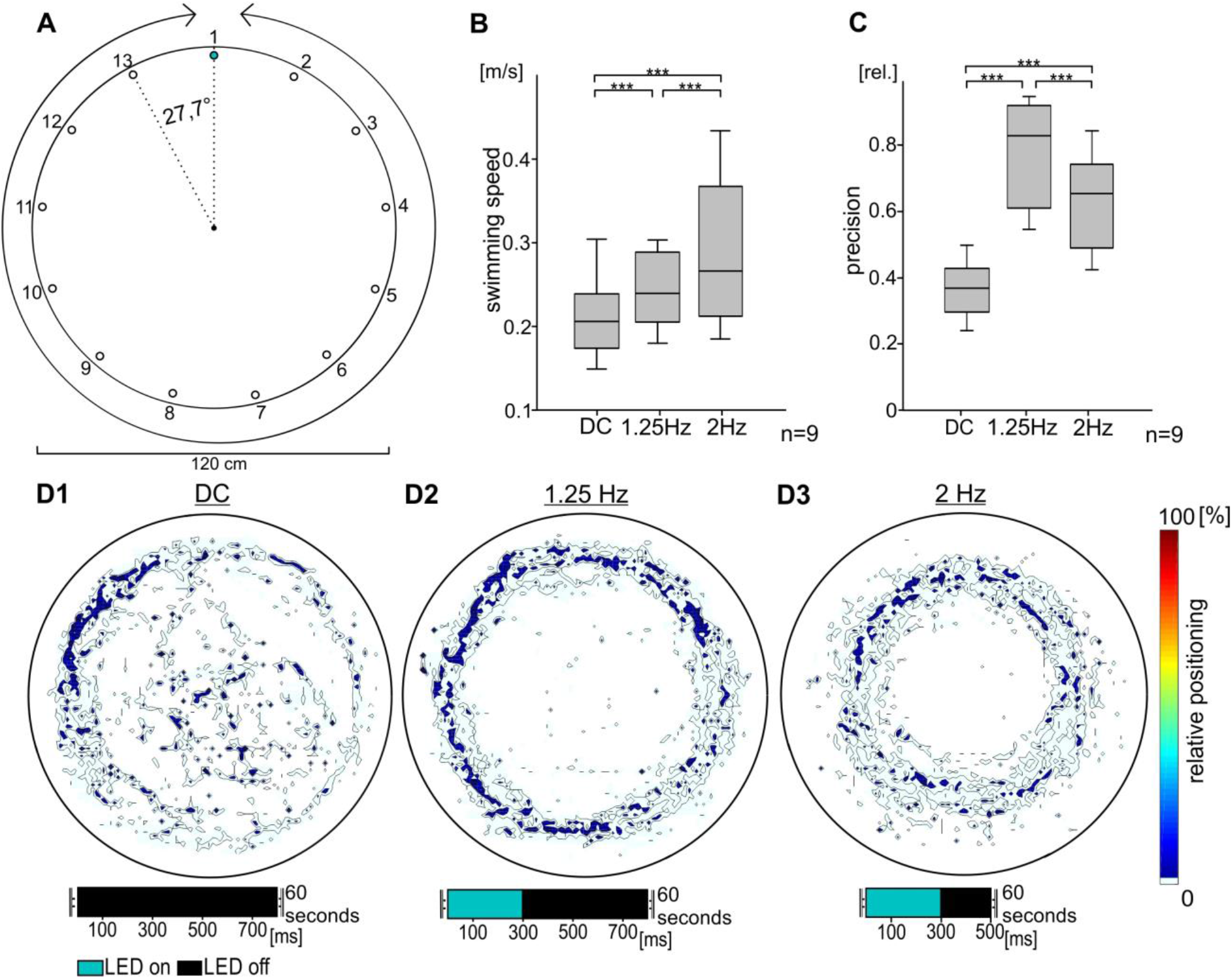
Motivation of flashlight fish (*A. katoptron*) to follow a moving light source. (A) Experimental setup with 13 wall mounted LEDs that were triggered consecutively counter- or clockwise. Intervals between 300 ms light emittance were 200 ms (2 Hz) or 500 ms (1.25 Hz) (travelling speed of light: 200 ms, 0.58 m/s; 500 ms, 0.36 m/s). Each fish was tested for 60 seconds in 5 trials. (B) Mean swimming speed of isolated *A. katoptron* (n=9) during control (DC, dark control), 200 ms off (2 Hz) and 500 ms off (1.25 Hz) times. (C) We estimated relative distance between specimen of *A. katoptron* (n=9) and the center of the tank according to the motivation of individuals to follow the moving light source. (D) Heat Maps indicate relative positioning of *A. katoptron* (n=9) during light stimulation (D2/D3) and control (D1, DC). Heat Maps are based on five trials for one isolated specimen. DC, dark control; *p < 0.05, **p < 0.01, ***p < 0.001.

Isolated specimens were following the counter- or clockwise rotating LED light to 75 % of the time without showing off-time-dependency (Fig. S3). A higher swimming speed of *A. katoptron* was observed for the 200 ms off-times (0.285 ± 0.013 m/s), representing faster moving LEDs, in comparison to the 500 ms off-times (0.246 ± 0.007 m/s) and the control without light stimulation (DC; 0.213 ± 0.008 m/s) (Fig. 3B). In contrast, the fish follows the rotating LEDs at 500 ms off-times closer and with higher precision (1.25 Hz; 0.771 ± 0.013; Fig. 3C, 3D2) in comparison to 200 ms off-times (2 Hz; 0.63 ± 0.023; Fig. 3 C, 3D3) and control (DC; 0.365 ± 0.013; Fig. 3C, 3D1). The results suggest that *A. katoptron* lose precision to follow artificial light organs at higher swimming speeds.

We next investigated the blinking behavior of several schools of *A. katoptron* in the ocean at a cave near Ambon and on a reef flat of Banda Island, Maluku, Indonesia. During the day the school of *A. katoptron* could be observed within the cave, while at sunset the school left the cave to approach the reef flat. We also observed a context dependent blink behavior and distinguished three different behavioral conditions, i.e. blinking behavior in the cave during the day, blinking behavior at the reef flat during the night and blinking behavior during avoidance triggered by a red diving torch. As also observed in the aquarium, blink frequencies increased from 1.96 Hz (cave, 117.69 ± 1.55 blink/min, Video S4), 3.33 Hz (reef flat, 199.71 ± 3.21 blink/min, Video S5) to 3.97 Hz (avoidance, 238.45 ± 4.79 blink/min, Video S6, Fig. 4A) with light organ occlusion ranging from 347.14 ± 10.8 ms (cave), 120.66 ± 2.39 ms (reef flat) to 68.65 ± 2.34 ms (avoidance, RM ANOVA: p < 0,001), while light organ exposure remained constant at around 230 ms (cave = 229.91 ± 3.05 ms, reef flat = 219.63 ± 4.79 ms, avoidance = 233.89 ± 6.53 ms, Fig. 4B). In addition, we found that the variation in blink frequencies is highest during avoidance behavior (Gaussian distribution; X (μ=3.97 Hz; σ²=3.062 Hz)) and low during daytime, while hiding in the caves (Gaussian distribution; X (μ=1.96 Hz; σ²=0.476 Hz)) (Fig. 4C). During avoidance behavior the relative nearest neighbor distance is reduced compared to reef flat schooling behavior from 2.03 ± 0.169 SL (n = 37) to 1.42 ± 0.09 SL (n = 46) and an increased group cohesion becomes obvious in the synchronized escape movements (Fig. 4D).

**Figure 4.**
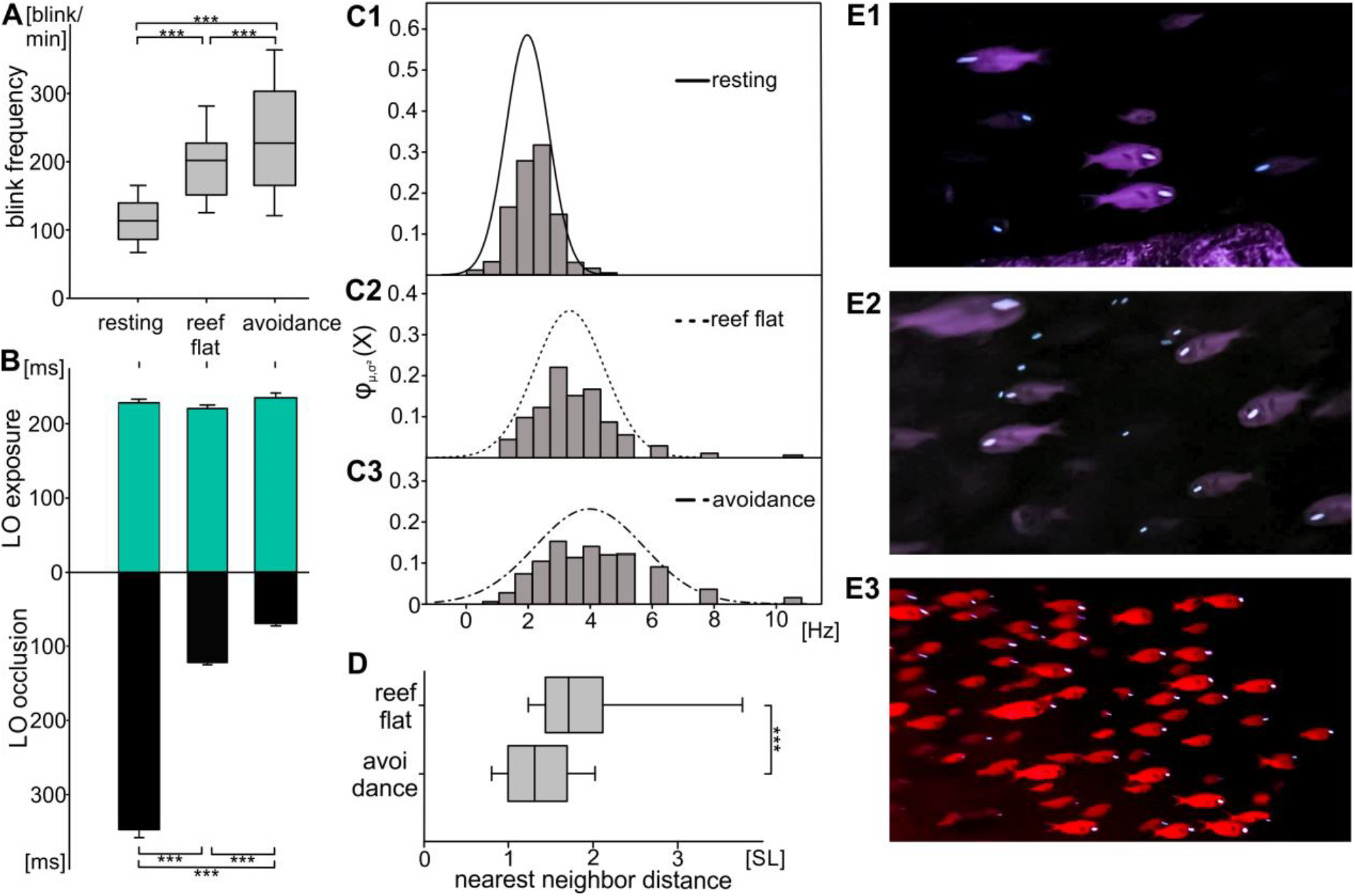
Analysis of the blinking behavior and nearest neighbor distance of schools of *A. katoptron* in Ambon, Maluku, Indonesia. The behavior of *A. katoptron* was analyzed for three conditions: in a cave during the day, during the night at the reef flat close to the cave and during avoidance behavior in the night. (A) Analysis of blink frequencies of *A. katoptron* in the cave, at the reef flat and during avoidance. Blink frequencies were calculated by analyzing alternating light organ exposition and occlusion (cave n=709; reef flat n=444 and avoidance n=478). (B) Mean light organ exposure and occlusion (± SEM) of *A. katoptron* in the cave (open n=823; closed n=761), at the reef flat (open n=502; closed n=445) and during avoidance (open n=516; closed n=478). Upper lines refer to stimulation as seen in (A). (C) Relative distribution of blink frequencies of *A. katoptron* observed while resting in the cave (C1), at the reef flat (C2) and during avoidance (C3). Bars represent histogram with bin size of 0.6 Hz. Distribution was fitted with normal (Gaussian) distribution (cave, X (μ=1.96, σ²=0.476); reef flat, X (μ=3,33, σ²=1.278); avoidance, X (μ=3.97, σ²=3.062)). (D) Analysis of the distance between specimen of *A. katoptron* on the reef flat and during avoidance. Screenshots of recordings were taken before (n=37) and during avoidance reaction (n=46) and analyzed. Avoidance was triggered by illumination of schooling *A. katoptron* with red diving torches. Distance is given as standard length (SL). (E) Example still images of the videos of *A. katoptron* schooling during day in the cave (E1), during the night on the reef flat (E2) and during avoidance behavior in the night (E3). Significance values are reported as: *p < 0.05, **p < 0.01, ***p < 0.001. Error bars indicate ± SEM

## Discussion

In this study we found that variation in blink frequencies of the bioluminescent splitfin flashlight fish *Anomalops katoptron* is used for intraspecific communication important for schooling behavior. Schools of *A. katoptron* can be observed at dark and moonless nights at the water surface in the Indo-Pacific. *A. katoptron* emit short bioluminescent light pulses using specialized light organs situated under the eye during schooling ^8,13^. These subocular light organs are densely packed with bioluminescent, symbiotic bacteria (*Candidatus photodesmus katoptron*), which continuously produce bioluminescent light ^14–16^. The fish disrupts light emission by a downward rotation of the light organ. Thus, exposure and occlusion of the light organ can produce specific blink frequencies ^45^. We found that adjustment of the blink frequencies of *A. katoptron* depends on variations within the occlusion and not the exposure of the light organ while schooling. Light organ exposure is comparable to previous laboratory (383 ms; ^8^) and field (166 ms; ^13^) studies. In comparison longer flash durations of 400 ms in *Lampanyctus niger* ^42^ and 1000-2000 ms in *Gazza minuta* ^44^ have been described in other bioluminescent fish.

Intraspecific recognition/communication is important to establish and maintain group structures ^46^. Species-specific signals like visual cues ^30,47^, motion ^48^, auditory ^49^ or electric signals ^37^ have been described to be involved in this process. Visual cues are important to detect position and movement of conspecifics ^31^ or predators ^50^ in fish and become crucial in species that live under dim/low light conditions such as *A. katoptron* ^7^. The bioluminescent light of *A. katoptron* is used for actively finding food and is most likely important for schooling behavior under dim light conditions and, therefore, for intraspecific communication ^8,13^. In our study we showed that *A. katoptron* follows moving LED light pulses and that the swimming speed is adjusted to the moving light. The speed of the moving LEDs (200 ms off-time; travel speed of light: 0.58 m/s) potentially exceeded the mean swimming speeds of *A. katoptron* (0,285 ± 0.013 m/s), since individuals could follow the moving LEDs at lower moving speed more precisely. Mean swimming speeds depend on various factors such as body size, tail beat frequency, scale types or hydrodynamic effects ^51^. The mean swimming speed of *A. katoptron* was estimated to 3,5 BL/s (body length per second), which correlates with other marine species ^24,52,53^.

We also analyzed the blinking behavior of *A. katoptron*. We found that for intraspecific recognition *A. katoptron* only uses information of the blinking light and not the body shape, since we did not detect differences in the blink behavior when we used LEDs or LEDs implanted within a fish dummy. We also found that higher light intensities of the LEDs induced lower blink frequencies of *A. katoptron*. One possibility is that higher intensity light is causing stronger behavioral responses, because the higher intensity light penetrates further through water and could be received as a closer schooling neighbor. We measured for the first time the maximal light intensity of light emitted by the light organ from *A. katoptron* which was at 0.27 μW at 510 nm wavelength. The retina of *A. katoptron* has also the maximal light sensitivity in this range ^19^. Light intensity could potentially represent fitness levels of individuals as *A. katoptron* tend to loose luminescence due to starvation ^54^. Other fish species prefer to shoal with healthy conspecifics ^55,56^. Schooling fish tend to show consistency in their appearance (confusion effect) ^29,57^ and often do not show a sexual dimorphism including flashlight fish (but also see pony fish *Gazza minuta*) ^44,58^.

The most important result of our study is that blink frequencies adjusted by light organ occlusion determine nearest neighbor distance. We suggest that light organ exposure and occlusion are alternating signals for attraction and repulsion in defining nearest neighbor distance in schooling *A. katoptron*. Nearest neighbor distance is a key factor in schooling fish and determines group cohesion ^59^. The shape of a school is the integration of individual responses on surrounding ecological factors ^46^. Thereby intraspecific signals such as bioluminescent blinks in flashlight fish need to be included. In ponyfish (Leiognathidae) luminescent flashes have been proposed to function in spacing between individuals and keeping the school together ^60^. Here we present a mechanism that potentially drives the opposing forces of attraction and repulsion in bioluminescent fish.

To gain an understanding of how blinking behavior is used for intraspecific communication in the field, we analyzed the blinking behavior of schools of *A. katoptron* in a cave during the day and at the entrance of the cave during the night in Ambon, Malukku, Indonesia using infrared recordings. We found that the blink frequency decreased during the day in comparison to blink behavior at night. Blinking behavior increased when fish were illuminated with a red torch, which caused an avoidance behavior and a reduction in the nearest neighbor distance. This became obvious by a change from a broad to a dense school formation. Increased blink frequencies seem to be correlated with stress reactions, since we observed increased blink frequencies when *A. katoptron* were isolated in the laboratory. Increase in blink frequencies are also known from *Photoblepharon steinitzii*, when artificial intruders have been introduced into their territory ^12^.

In conclusion, our study shows that *Anomalops katoptron* uses intraspecific, bioluminescent blink signals for communication of nearest neighbor distance important for group cohesion during schooling.

## Methods

### Recordings in the Laboratory

#### Maintenance of *A. katoptron*

A group of splitfin flashlight fish *A. katoptron* was kept in a reef tank (600 l; 135 cm length × 66 cm depth × 70 cm height). All specimens were obtained from a commercial wholesaler (De Jong Marinelife, Netherlands) and captured at the Cebu Islands (Philippines). For at least six weeks prior to the experiments *A. katoptron* were kept in the reef tank (temperature: 26°-27°C; salinity: 36 ‰; 12 h day and night cycle). The housing tank (600 l; 135 cm length × 66 cm depth × 70 cm height) was connected to an additional filter sump containing phosphate absorber, activated carbon, protein skimmer and an UV-sterilizer. The specimens were fed once a day with defrosted zooplankton (mysid shrimps), fish/lobster eggs and fine minced, defrosted salmon. Feeding occurs under dim red light to obtain visual observation on fitness levels of individuals. Information on age is missing because all individuals were wild collected imports. No visible differences between females and males were observed. Individuals were identified by size, slight differences in pigmentation and intensity of light organs.

#### Artificial light organs and fish dummies

A fish dummy with artificial light organs was made of black silicon (food safe silicon MM720 FG; Silikonfabrik; Germany). The shape of the dummy was modelled based on several photographs and had a total length (TL) of 101 mm. At the anterior-ventral side an oval shaped opening was cut out of the dummy. The cutout was equipped with a LED to imitate the light organs of *A. katoptron*. The LED was connected to an Arduino microcontroller (Arduino Mega 2560; Arduino; Italy). Resistors between LED and microcontroller were set to an output flow of 1 mA. The LED was waterproof glued (2-K epoxy glue; UHU; Germany) in an acrylic glass tube (length 15 mm; external diameter 7 mm) painted with flat white acrylic paint (Revell; Germany) to diffuse the LED light. The acrylic glass inlet was mounted in the fish dummy (artificial light organ length: 10 mm; height: 7 mm). The LED (Nichia 3mm LED cyan 14.720mcd; Winger; Germany) had a peak wavelength at 500 nm and was adjusted to the mean light emittance of 0.23 μW/nm of *A. katoptron’s* light organs (Fig. S2). Intensities of light organs (n=5) and LEDs were measured with a spectrometer (Ocean Optics; Flame; United States).

The microcontroller was set to control artificial light organs in relation to on- and off-times. The control software was written with Matlab (Matlab 2015r) and the open source Arduino software (Arduino 1.8.10). LED light intensities were adjusted by using a pulse width modulation (PWM).

Recordings in the experimental tank and arena experiments (see below) were made with an infrared (IR) sensitive camcorder (Sony HDR-CX 730; 6.3 mm CMOS-Sensor, 24.1 megapixel, video resolution 1920 × 1080 pix, 50 fps) mounted on a custom made aluminum stand. Video files were converted to audio video interleave-format (.avi) with a resolution of 1080 × 720 pix and 25 fps using Adobe Premiere Elements 15 (Adobe; United States).

#### Blink frequencies (equal LED on- and off-times)

The recording tank was divided in the middle with a grey PVC plate. Specimens could switch sides through a lockable slide door (20 × 20 cm). One of the sides contained daytime shelters made from clay tiles whilst the other half was blank except for a flow pump (EcoDrift 4.2; Aqua Medic; Germany). Specimen of *A. katoptron* (n=5) were isolated on the blank side (60 cm × 60 cm × 60 cm) of the experimental tank and habituated for five minutes prior to the experiment.

The fish dummy was placed in the middle of the recording tank. Each light stimulus was presented for a duration of five minutes. Every stimulus presentation was repeated five times. Here we chose equal distributed on- and off times in LED timing with 0.5 Hz (1 s on- and 1 s off-time), 1 Hz (0.5 s on- and 0.5 s off-time) and 2 Hz (0.25 s on- and 0.25 s off-time). Previous laboratory experiments showed a nearly equal distribution of light organ exposure and occlusion times while swimming in a group ^8^. We performed a control experiment with turned off artificial light organs (DC, dark control) implemented in the dummy. The camera was mounted on a tripod in front of the tank. Two IR-lights each consisting of five high power LEDs with 860 nm peak wavelength (WEPIR1-S1 IR Power 1 W, Winger Electronics GmbH, Germany) were placed 10 cm above the tank.

In a second experiment, we analyzed the role of dummy (fish) shape and isolated light organ dummies on the behavior of *A. katoptron* (n=5; same individuals used in the first experiment). Therefore, an isolated light organ dummy (LED as described above) was used during stimulation. We chose a light stimulation protocol of 1 Hz (0.5 s on- and 0.5 s off-times) because this stimulation had the strongest effect on blink frequencies of isolated specimen. In the next step we analyzed differences in blink frequencies for two specimens with intact light organs as well as one specimen with intact and one with non-glowing light organs to test orientation of *A. katoptron* towards light organs of conspecifics (Fig. S1). In this case, we performed a frame by frame analysis (video analysis software; Vidana 1.0) of distances between individuals. All stimuli were presented for five minutes in a pseudo-randomized order. Five repetitions were performed for each specimen.

Blink frequencies (reported in blink/min) and light organ exposure -/occlusion-times (reported in ms) were analyzed frame by frame using Solomon Coder (Version 19.08.02). Mean values of blink frequencies and light organ exposure-/occlusion-times were analyzed with Excel (Excel 2016). Successive exposure and occlusion events were summarized as blink event.

Trajectories were analyzed with the video analysis software Vidana 1.0. Two rectangles of interest (ROI) were defined to analyze the swimming profiles in *A. katoptron*. As individuals could switch between the two sides of the tank amongst experiments, we defined the areas where occurrence was most likely. The area around the closed door was declared as “exit area”. The area around the dummy placed in the middle was defined as “center”.

#### Arena experiment 1: Nearest Neighbor Distance

Large-scale swimming profiles during presentation of a fish dummy with artificial light organs were analyzed in a circular arena with 120 cm diameter (Winipet Dogpool; China). Seawater from the housing tank was used to ensure equal parameters in water chemistry e.g. carbon hardness, nitrate and pH values. The arena was filled with approximately 170 l seawater (15 cm water level). Single specimen of *A. katoptron* (n=10) were transferred to the arena using a hand net (12.5 cm × 10 cm; Sera; Germany). Prior to the experiments fish were habituated for five minutes in the arena tank. A fish dummy with artificial light organs (as described above) was placed 7.5 cm over the tank bottom in the center of the arena. In this experiment artificial light organs were constantly glowing up for 300 ms whereas off-times changed. The occlusion of artificial light organs was adjusted to 200 ms (2 Hz stimulation) or 500 ms (1.25 Hz stimulation) but consistent during one trial. We additionally performed a control experiment without light emitted by the dummy (DC, dark control). Stimuli were randomly presented for 30 seconds with six repetitions. Videos were recorded using an infrared (IR) sensitive Sony HDR-CX730E camcorder (1920 × 1080 pix; 50 fps) mounted above the arena on a custom made stand. Two IR-lights each consisting of five high power LEDs (WEPIR1-S1 IR Power 1 W, Winger Electronics GmbH, Germany) were placed besides the arena mounted on custom made holding devices. Tracking profiles of *A. katoptron* were analyzed using the video analysis software Vidana 1.0. Heat maps were generated in Matlab (Matlab R2015b). Here we summarized equal positions of standardized tracking profiles to estimate relative occurrences of *A. katoptron*.

#### Arena experiment 2: Swimming Speed

To validate the following behavior and maximum swimming speeds of *A. katoptron* we established an array of LEDs that were rotated consecutively to simulate a moving light organ. In this experiment, 13 LEDs were wall-mounted in an equal distributed distance (specifications circular arena see above). Angle between LEDs was set to 27.69°. The LEDs were placed on a water level of 7.5 cm. On-times of LEDs was permanently set to 300 ms while interval among the light onset between two LEDs was changed. During one trial intervals between two LEDs were set to 200 ms or 500 ms. LEDs were triggered clockwise or counter clockwise in a pseudo randomized order. A dark control (DC) experiment without light stimulation was performed to avoid potential orientation cues from the periphery of the experimental arena. Handling of *A. katoptron* as described under Arena Experiment 1.

Experiments in single specimens of *A. katoptron* (n=9) were started after five minutes habituation time in the arena. Each stimulus was presented for 60 s. Specimens were tested five times for each stimulus. Movement profiles, swimming speed and radius of *A. katoptron* were analyzed using the video analysis software Vidana 1.0. Relative movement directions were estimated with Solomon Coder (Version 19.08.02). We estimated the precision of *A. katoptron* to follow moving light sources on a defined radius (distance between individuals and center of the tank). For each stimulation (1.25 Hz, 2 Hz & DC) we calculated the probability of individuals to move with the direction of light (Fig. S3). During dark control (DC) isolated individuals were moving clockwise (0.41 ± 0.034), counterclockwise (0.44 ± 0.034) or without a defined movement direction, declared as other (0.15 ± 0.001). Isolated specimen were following the counter- or clockwise rotating LED light to 0.724 ± 0.034 (200 ms off-times) and 0.78 ± 0.031 (500 ms off-times). Subsequently we multiplied the probability to follow the rotating light or the highest value in case of the dark control (DC) with the radius to estimate the precision.

#### Field Recordings

Field recordings were made alongside two different Islands in the Banda Sea (Indonesia). Several schools of *A. katoptron* were observed via snorkeling on the shallow reef flats of Pulau Gunung Api, Banda Islands (4°30’20.2”S 129°52’49.7”E). Recordings on the Banda Islands were made after sunset on 1^st^-4^th^ of March 2019 prior new moon (7^th^ of March 2019) and the 26^th^ of March 2019 (five days after full moon). Recordings on the Banda Islands were made before moonrise. Schools of *A. katoptron* occur from deeper water (> 60 m; pers. obs.) or caves during dark and moonless nights on the shallow reef flats of Gunung Api. The observation site in Ambon (3°44’54.5”S 128°12’43.3”E) was quite different and recordings made while scuba diving. Schools were hiding throughout the day in a large cave (main chamber dimensions approximately 10 × 5 × 6 m) with many small crevices that were not accessible. The cave entrance was in approximately 6 m depth beneath the water surface depending on the tide. Field recordings in Ambon were made between 19^th^-20^th^ of March 2019 before full moon (21^st^ of March) and on 17. April 2019 before full moon (19^th^ of April 2019). During the day, recordings were made in the cave and continued while sunset when schools of *A. katoptron* emerged through the cave exit. After several minutes schools accumulated in front of the cave where overhanging rock casts a shadow of the moonlight. This was leading to a restricted area of movement. We defined three different recording conditions to analyze the behavior in *A. katoptron*: 1. “resting” (recordings in the cave during day without illumination); 2. “schooling” (outside cave or on reef flat during night without illumination) and 3. “avoidance” (avoidance elicited by red diving torch during night).

Video recordings were made with a modified camera (Canon Powershot G1X Mark 2; APS-C-Sensor; 24 megapixel; video resolution: 1920 × 1080 pix; 30 fps). The infrared filter in front of the camera sensor was removed to obtain infrared sensitivity. The camera was placed in an underwater housing (Canon WP-DC53). Two custom made underwater infrared lights mounted on both sides of the underwater housing were used two illuminate schools of *A. katoptron* in the cave and open water. Each IR-light consisted of five high power IR-LED with 860 nm peak wavelength (WEPIR1-S1 IR Power 1 W, Winger Electronics GmbH, Germany).

A LED diving torch with red light (300 lumen red light; 634 nm peak wavelength; Codylight 1500; Codygear; Germany) was switched on while the school was swimming outside the cave or on the reef flat to elicit avoidance reactions. “Avoidance” was triggered pseudorandomized when specimen were within a range of approximately 1.5 m to ensure sufficient illumination with IR-lights. The red light was switched on until the school disappeared from view. After *A. katoptron* gathered outside the cave a minimum of two minutes was waited before red torches were repeatedly turned on.

We recorded n=5 video sequences (709 blink events in 326 seconds) for “resting” in the cave, n=8 video sequences (444 blink events in 272 seconds) during “schooling” on the reef flat and n=5 video sequences (478 blink events in 40 seconds) in case of “avoidance”.

Relative distances between school members were estimated via ImageJ (ImageJ 1.50i; National Institute of Health). We compared single screenshots taken from video sequences of schooling *A. katoptron* without (n=37) and with illumination with red torches (n=46). We defined relative length (SL) of at least one individual as reference to estimate the relative distance between members of the school. We chose distances between individuals that seemed to be neighbors as two-dimensional recording could not provide a distinct spatial distribution (see also Fig. S4).

Blink frequencies were analyzed using the video analysis software Vidana 1.0. Specimens of *A. katoptron* were marked after the first occurrence in the video sequence and the behavior was analyzed until the specimen disappeared in the recording sequence. Exposure and Occlusion of light organs was analyzed frame by frame per individual occurrence. Mean values were summarized for all analyzed parameters. Blink frequencies were estimated based on pairs of light organ exposure and occlusion times. We created a Gaussian distribution (Fig. 4) using the internal SigmaPlot function (SigmaPlot 12.0) to show the distribution of blink frequencies during three situations in the field (“resting”, “schooling” & “avoidance”). Additionally, we created histograms with the internal Matlab function (Matlab R2015b). Here we chose a bin size of 0,6 Hz.

#### Statistical Analysis

SigmaPlot 12.0 was used to evaluate statistical differences between test groups. Differences in blink frequencies, exposure and occlusion times of light organs, distance between individuals, swimming speed and spatial distribution were compared using a repeated measurement one-way ANOVA and Holm-Sidak post hoc analysis. All values are reported as mean ± SEM (standard error of mean). Statistical significant values are reported as: * p ≤ 0.05, ** p ≤ 0.01; *** p ≤ 0.001.

## Supplemental Figures

**Figure S1.**
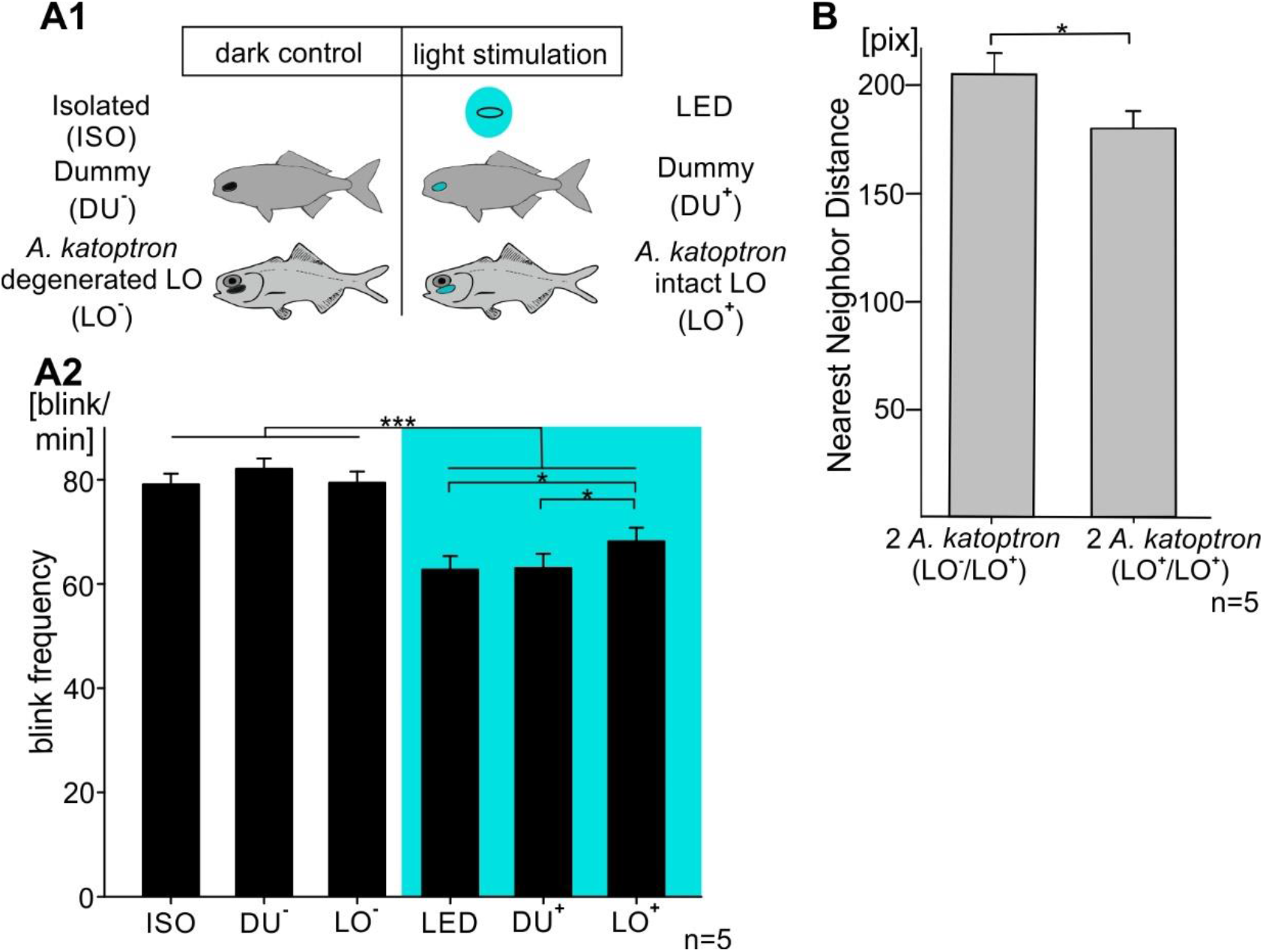
Blink behavior of *A. katoptron* during exposure to different artificial light stimuli and their orientation towards conspecifics. (A1) Natural and artificial light stimulation used in the experiments to investigate the blink behavior of *A. kataptron*. Two different types (light stimulation or dark controls) were presented to isolated individuals, i.e. an isolated LED and a fish dummy, equipped with a LED at the position of the light organ. Artificial lights had the same size, intensities and emitted 1 Hz light pulses (equally distributed LED on- and off- times). Experiments were repeated 5 times with five individuals independently and values are given as mean (± SEM). The behavioral responses to the artificial lights were compared to responses to *A. kataptron* with an intact light organ (LO^**+**^) and a degenerated light organ (LO^**−**^). (A2) Blink frequencies of isolated *A. katoptron* during exposure to dark controls (left, white background) and light stimulation (right, blue background). Blink frequencies were reduced in the presence of light-stimuli. LED and the fish dummy light stimuli reduced the blink frequency more than the conspecifics. (B) *A. katoptron* show a closer mean (± SEM) orientation towards its neighbors when both specimen display intact light organs (LO^**+**^/LO^**+**^). Statistical significance was evaluated with RM ANOVA. Significance values are reported as: *p < 0.05, **p < 0.01, ***p < 0.001. Error bars indicate SEM.

**Figure S2.**
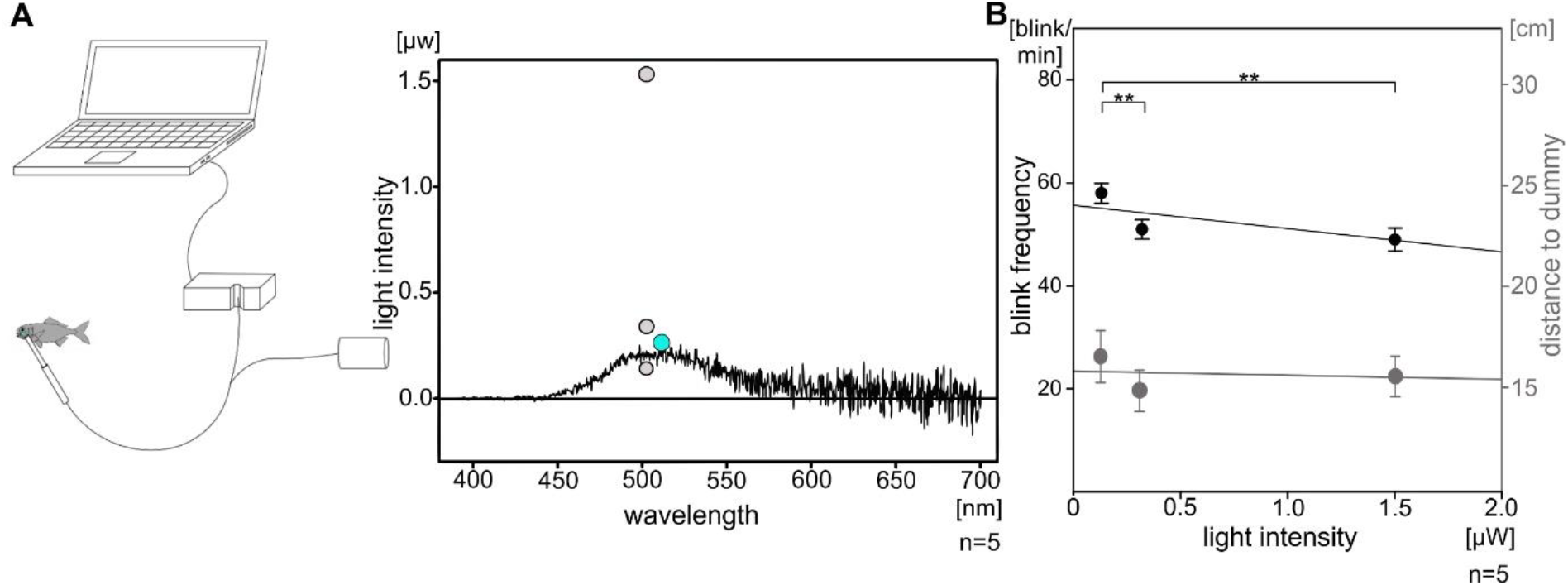
Spectrometric measurements of the light intensities emitted by light organs of *A. katoptron* in comparison to LEDs and the dependency of blinking behavior on different light intensities. (A) Spectrometric measurement of the light organ (n=5) intensity of *A. kataptron* in comparison to LED light. Diagram of the experimental setup for the spectrometric measurements (left). Intensities were measured with a spectrometer (Flame S-UV-VIS-ES, Ocean Optics, USA). The spectroscopic probe was placed in front of the light organs of fixed individuals. Each light organ was measured five times and mean intensity averaged. Example trace of the spectrometric measurement of the light intensities emitted by the light organs measured in the range between 400 – 700 nm wavelength (right). Gray dots indicate three different LED intensities of 0.133, 0.328 & 1.523 μW at 504 nm wavelength, which were presented to isolated flashlight fish (*A. katoptron*) to investigate impact on blink frequency. The green dot indicates the maximum intensity observed in *A. katoptron* (0.27 μW at 510 nm wavelength). (B) Blink frequency responses of *A. katoptron* and distance to dummy triggered by the three distinct LED intensities detected at 504 nm wavelengths as shown in A. Experiments were repeated 5 times independently and values are given as mean (± SEM). Blink frequencies of *A. katoptron* were decreased with increasing intensities of the LED. Statistical significance was evaluated with RM ANOVA. Significance values reported as: **p < 0.01. Error bars indicate ± SEM.

**Figure S3.**
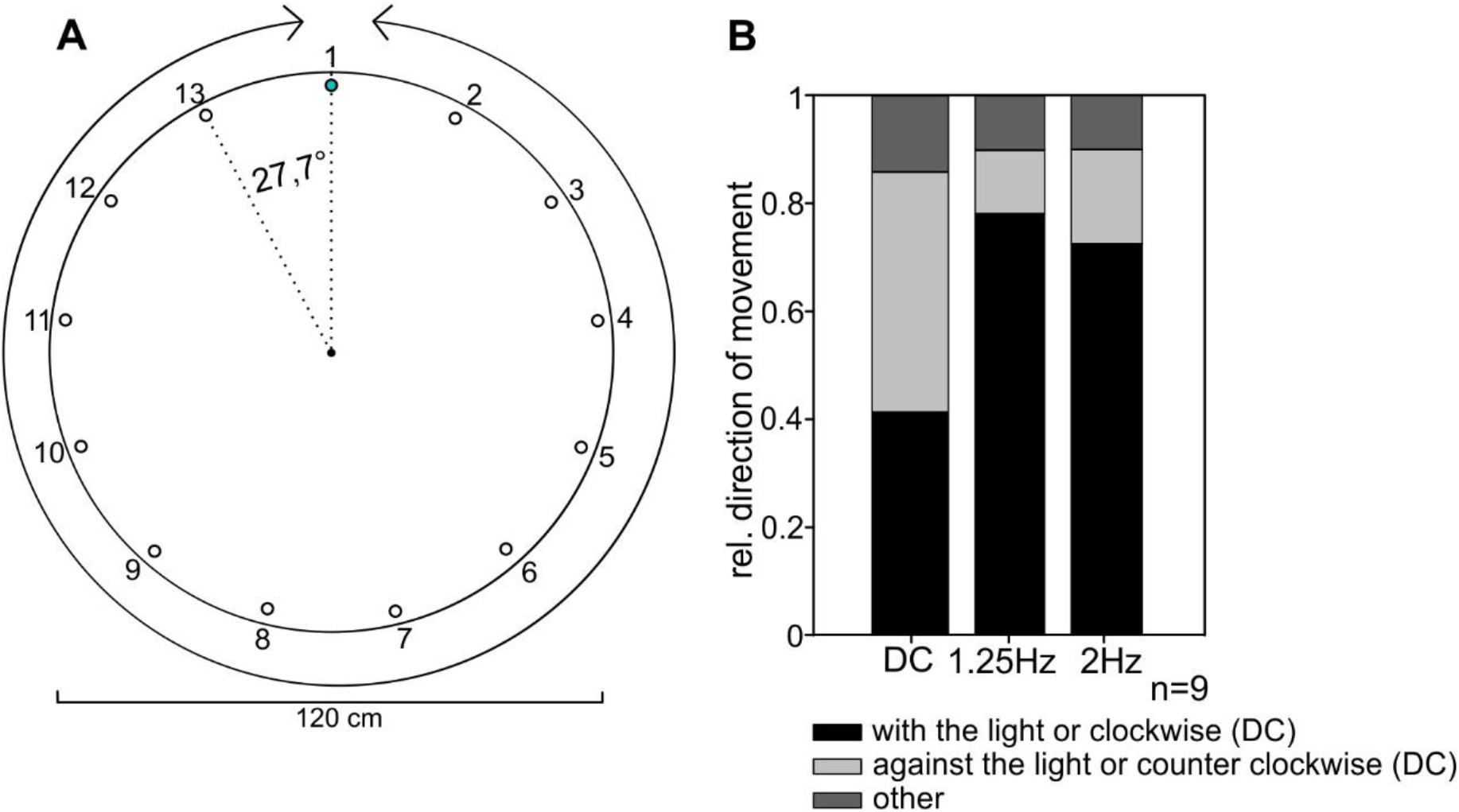
*A. katoptron* follow moving light stimuli. (A) Experimental setup with 13 wall mounted LEDs that were triggered consecutively counter- or clockwise. Intervals between 300 ms light emittance were 200 or 500 ms (travelling speed of light: 200 ms, 0.58 m/s; 500 ms, 0.36 m/s). Additionally, we performed a control without light stimulation (DC, dark control). Each fish was tested for 60 seconds. (B) Relative direction of *A. katoptron* following artificial light sources. Flashlight fish show a high motivation to follow the direction of light (1.25 & 2 Hz). In the control experiment (DC) the fish swims equally into both (counter clockwise or clockwise) directions.

**Figure S4.**
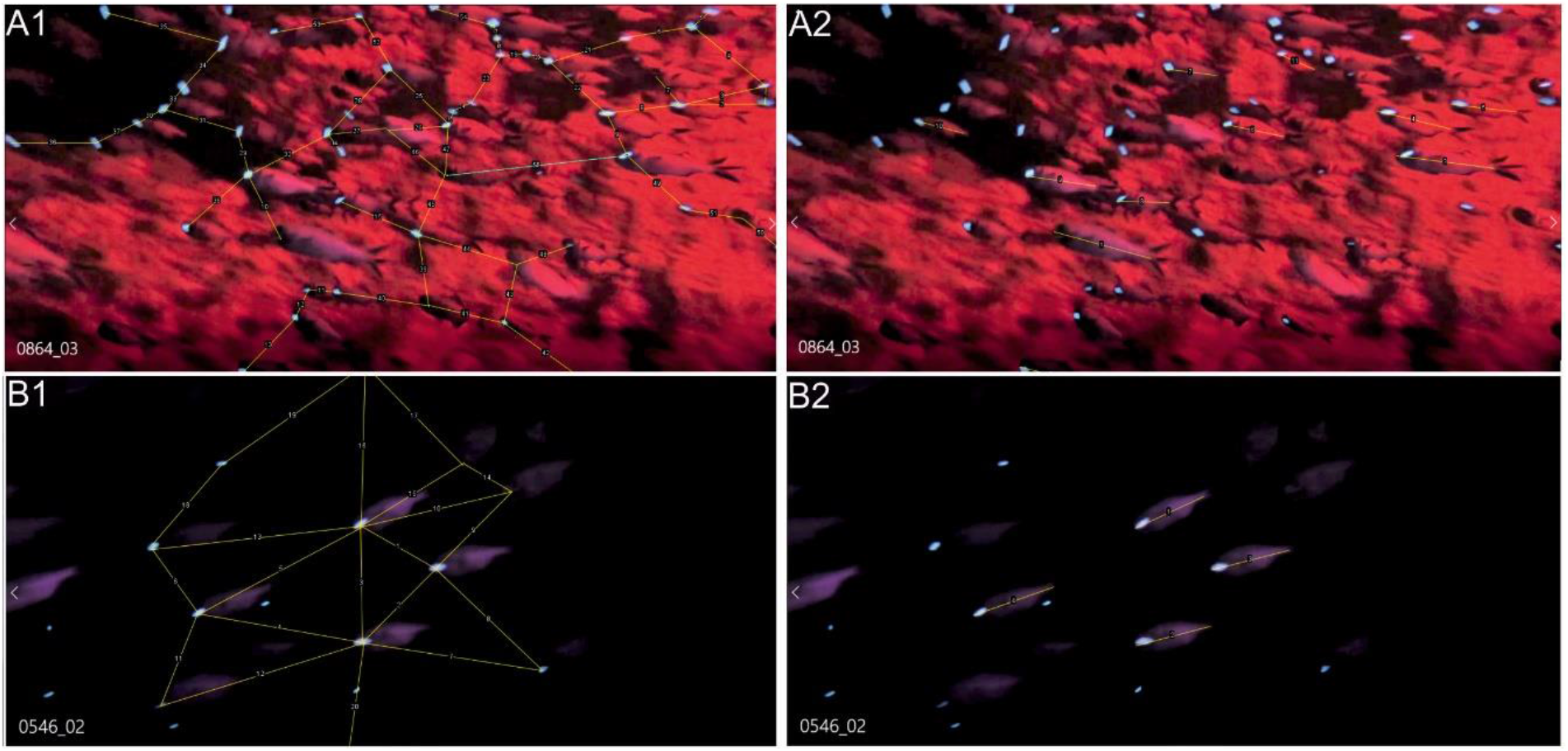
Analyzing the nearest neighbor distance in schools of *A. katoptron* (Ambon, Maluku, Indonesia). (A) Flashlight fish *A. katoptron* were illuminated with diving torches (300 lumen red light; Codylight 1500; Germany) to trigger avoidance reactions. For every screenshot, we estimated the fish standard length (SL) as reference (A2) We connected light organs (A1) of individuals that seemed to be neighbors to determine their distances. 46 screenshots were analyzed. (B) Groups of *A. katoptron* while schooling on the reef flat were illuminated with IR-torches and recorded with an infrared camera. Networks connect light organs of potential neighbors (B1) and standard length (SL) was estimated as reference (B2). 37 screenshots were analyzed.

## Notes

### Competing Interest Statement

The authors have declared no competing interest.

